# Direct-view oblique plane microscopy

**DOI:** 10.1101/2025.02.11.635467

**Authors:** Jacob R. Lamb, Miguel Cardoso Mestre, Madeline Lancaster, James D. Manton

## Abstract

The ability to rapidly image mesoscopic samples is critically important for many areas of biological research. Owing to its high throughput and gentle, volumetric imaging nature, light-sheet fluorescence microscopy (LSFM) is an attractive modality to image such samples. However, the orthogonal dual-objective geometry of LSFM makes sample mounting challenging. Oblique plane microscopy (OPM) circumvents these issues by achieving light-sheet imaging through a single objective near the sample, with a further two objectives acting to image a tilted, oblique plane in the specimen. However, at low magnification, as required for mesoscopic imaging, conventional OPM systems suffer from a complicated and inefficient light path and/or a limited numerical aperture. Here we present a simple method for mesoscopic OPM that enables efficient light collection at any numerical aperture. By placing the camera directly in the remote space, the oblique plane can be imaged without the need for a third microscope system. Using a commercially available camera with 1.4 µm pixels, we demonstrate imaging over a 5.3 × 3 × 2.6 mm^3^ field of view with a lateral resolution of ∼2 µm and axial resolution of ∼22 µm. Our technique is then used to image a 4×-expanded brain organoid.

## 1 Introduction

With the aim of investigating biological processes in models that better recapitulate the true in vivo micro-environment, many studies are turning to large 3D samples such as organoids [1, 2] and tissue culture models [3]. Furthermore, developments in tissue clearing [4] and expansion microscopy [5] now make it possible to render these large samples optically transparent, enabling the use of high-resolution optical imaging techniques. Consequently, there is increasing demand for 3D fluorescence imaging methods suited to mesoscopic scale samples.

Among the available techniques, light-sheet fluorescence microscopy (LSFM) is the most widely applied to mesoscopic samples due to its inherently high throughput and gentle volumetric image approach. Here, two orthogonal objectives are used: one generates a sheet of excitation light, while the other collects the fluorescence. This axially confined excitation allows for rapid generation of high-contrast, optically sectioned images of large samples. However, the need for two orthogonally mounted objectives imposes geometric constraints on sample mounting. To address this, variants of LSFM have been developed to simplify sample mounting. Of these methods, a configuration, known as oblique plane microscopy (OPM), has gained much popularity due to its ability to perform both light-sheet excitation and fluorescence detection through a single objective lens near the sample [6]. This allows for high quality volumetric imaging of samples mounted for inverted epifluorescence microscopy.

Central to the concept of OPM is the ability to create an aberration-free 3D image of the sample through a technique called remote refocusing [7, 8]. Fundamentally, remote refocusing consists of two microscopes that are placed back-to-back, so that the magnified image produced by the first microscope is demagnified by the second. Provided that the ray angles are conserved between the object and remote space, the demagnified image is free from the spherical aberrations that are normally present away from the native focal plane. This allows for an oblique light-sheet to be generated in the sample and imaged by a third, tilted microscope in the remote space. However, due to the limited solid angle of the light cone leaving the second objective, a greater proportion of the light is not collected as the third objective becomes more inclined. This has the combined effect of reducing the spatial resolution, resolution isotropy and photon efficiency. To mitigate this, OPM is typically limited to high magnification, high numerical aperture (NA) implementations where the large solid angles ensure sufficient light capture. As the magnification and NA of the system decrease, light loss increases until total loss of light occurs at 0.5 NA (assuming the use of three identical air objectives), making conventional OPM unsuitable for mesoscopic applications.

Despite this, OPMs operating at NAs below 0.5 have been reported. These methods either redirect light toward the third objective using optical components such as diffraction gratings [9, 10] or rigid fibre plates [11], or they reduce the relative angle between the second and third objectives. This can be done by either forgoing the aberration-free condition for remote refocusing [12] or creating shallower light-sheet angles than what the low NA primary objective allows. These shallower angles can be achieved through diffraction gratings [13], mirroring [10], or a separate excitation lens [14, 15]. Such methods also increase axial resolution and optical sectioning strength. Despite enabling low NA OPM, current solutions suffer significant drawbacks. For example, diffraction gratings split light into multiple diffraction orders which significantly reduces photon efficiency, rigid fibre plates are expensive, and forgoing aberration free remote refocusing reintroduces axially dependent spherical aberrations which reduce image quality.

To address these limitations, we present an alternative approach to mesoscopic OPM that collects the full collection cone of the primary objective lens at any NA, whilst also increasing photon efficiency. In our method, termed directview OPM (DvOPM), the camera is placed in the remote space where it directly captures the inclined plane. This approach removes the third microscope system used in conventional OPM, thus eliminating any need for optical components to redirect the light in the remote space. Using a commercially available backside-illuminated (BSI) CMOS camera with 1.4 µm pixels, an implementation of our method achieves a lateral resolution of 2 microns over a FOV of 5.3 × 2.8 mm in the *xz* plane. In a proof-of-concept experiment, we demonstrate the imaging capability of the system by imaging a 4×-expanded brain organoid.

## 2 Methods

The optical layout for direct-view OPM is summarised in Fig. 1. Currently, the system is optimised for imaging watery samples, thus the magnification of the remote refocus system should be matched to 1.33 [8, 16]. To minimise complexity, this magnification is achieved using only commercially available lenses. The pupil plane of an Olympus 4× 0.16 NA objective (UPLXAPO4X), O1, is mapped onto the pupil plane of a Nikon 4× 0.2 NA objective (CFI Plan Apochromat Lambda D 4X), O2, using a 4 *f* relay. This relay consists of an *f =* 200 mm tube lens (Thorlabs TTL200A), TL1, and an *f =* 165 mm tube lens (Thorlabs TTL165A), TL2. This combination gives a total magnification between the sample and remote space of 1.347×, which is within 1.3 % of the desired magnification.

**Figure 1:**
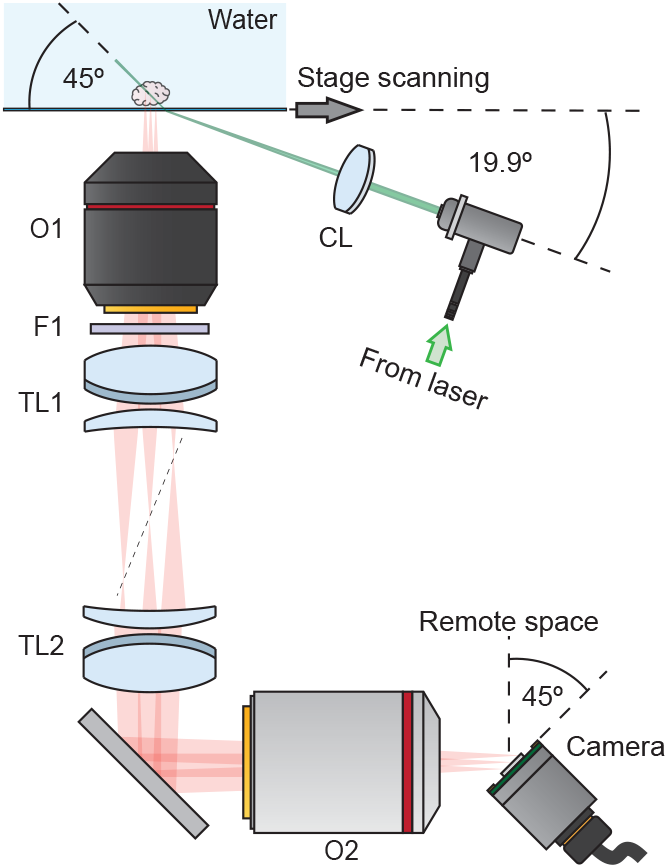
Schematic for direct-view OPM. A cylindrical lens, CL, is used to create a 45° light sheet in the sample. Fluorescence is collected by the primary objective, O1, and a remote image is formed after passing through a filter, F1, two tube lenses, TL1 and TL2, and a secondary objective, O2. A camera, at a matching angle to the light sheet, detects the inclined image in the remote space.

Due to the small angle of the primary objective’s light cone (18.4°), the minimum sheet angle that it can create is 80.8°. Therefore, to create a shallower sheet, improving both axial resolution and detection efficiency, we decouple excitation and emission by using a second lens to generate the light sheet. Excitation light is generated using a fibre coupled laser (FISBA READYBeam Bio1) and is collimated by a reflective fibre collimator (Thorlabs, RC08APC-P01). The excitation light is then focused into the sample using an *f =* 100 mm cylindrical lens (ACY254-100-A), CL. To achieve a sheet angle of 45° in the sample, the optical axis for the excitation light in air is at an angle of 19.9° to the coverglass. As the NA of the light sheet is very small (∼0.02), aberrations from traversing the coverglass at an angle are negligible.

The remote space fluorescence image is directly detected using a small camera (Ximea MU196MR-ON), with the sensor cover glass removed to avoid aberrations (Fig. S2 and S3), angled at 45° and brought to be coplanar with the light sheet plane. This camera uses a 7.1 × 5.3 mm^2^ sensor with 1.4 µm pixels (onsemi AR2020) and a maximum quantum efficiency of 91 %. Accounting for the magnification of the remote refocus system, this gives an effective pixel size in the sample of 1.04 µm, giving a sampling-limited resolution of 2.08 µm over a FOV of 5.3 × 2.8 mm^2^ in the *xz* plane.

Full details of organoid sample preparation are given in Supplementary Note 1. Briefly, 2000-starting-cell brain organoids were grown according to standard protocols. After 12 days of growth, organoids were fixed using 4 % paraformaldehyde at room temperature and expanded using the U-ExM protocol [17]. Nuclei were stained using Hoescht 33342.

## 3 Results

Though the method we present here increases photon throughput significantly over alternative approaches, it is important to note that increasing the incident angle of light onto the camera decreases the detection efficiency. To quantify this effect, we measured the variation in camera detection efficiency as function of incident ray angle for both S- and P-polarised light. This was achieved by measuring the total detected counts (after background subtraction) on the camera when illuminated with a collimated 532 nm laser beam at angles from 0 to 55°. Figure 2a presents these results with efficiency at each angle normalised by dividing the total measured counts by that measured at 0°. At an incident angle of 45° the measured detection efficiency was 78.2 *±* 1.3 % and 75.4*±*0.3 % for S and P polarised light, respectively. Using linear interpolation, the average detection efficiency of photons in a 0.16 NA light cone, incident on the camera tilted at 45°, was calculated to range from 96.1 *±* 0.1 % to 54.4 *±* 9.9 %. Despite the reduction in camera efficiency, this still demonstrates a notable increase in overall detection in comparison to other mesoscopic OPMs with similar FOVs, which report photon efficiencies ranging from only 16 % [13] up to 48 % [11].

**Figure 2:**
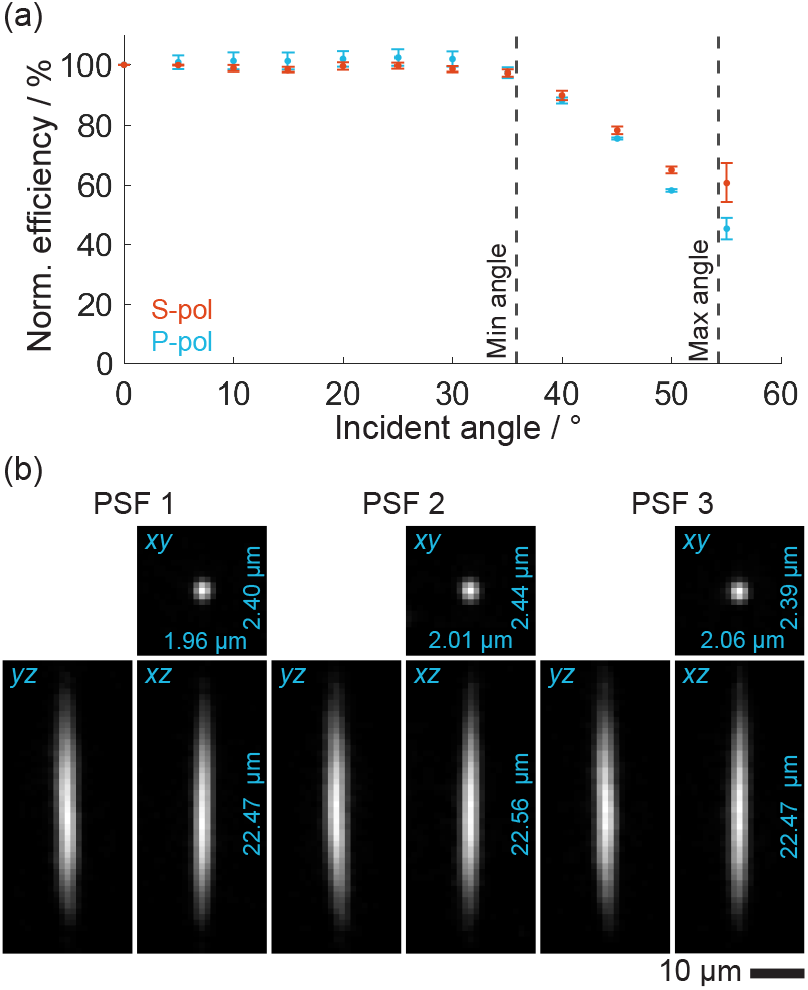
System characterisation. (a) Camera efficiency as a function of incident ray angle normalised to efficiency at 0°. The dotted lines indicate the minimum and maximum incident angle in a 0.16 NA light cone for a camera at 45°. (b) Exemplar beads used for spatial resolution measurement.

To characterise the spatial resolution of the system we imaged 200 nm diameter FluoSphere Yellow-Green beads dried onto a coverglass using a 0.02 NA lightsheet and a stage-scan slice interval of 1 µm. Spatial resolution was calculated by fitting a 3D Gaussian function to the deskewed and rotated data and extracting the full width at half maximum (FWHM) of the beads along each of the axis from the fitted standard deviation, *σ*, where 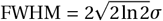. From an average of 12 beads, the measured spatial resolution is 2.05 *±* 0.06 µm × 2.42 *±* 0.02 µm × 22.31 *±* 0.35 µm. Figure 2b presents exemplar cross sections from three beads used to calculate the spatial resolution of the system.

To demonstrate the imaging capabilities of the system we apply the technique to imaging a 4×-expanded brain organoid. As can be seen in figure 3, direct-view OPM is capable of imaging the entire expanded organoid in a single volume whilst retaining sub-micron effective resolution. This allows for complex cellular organisation and structure to be rapidly visualised.

**Figure 3:**
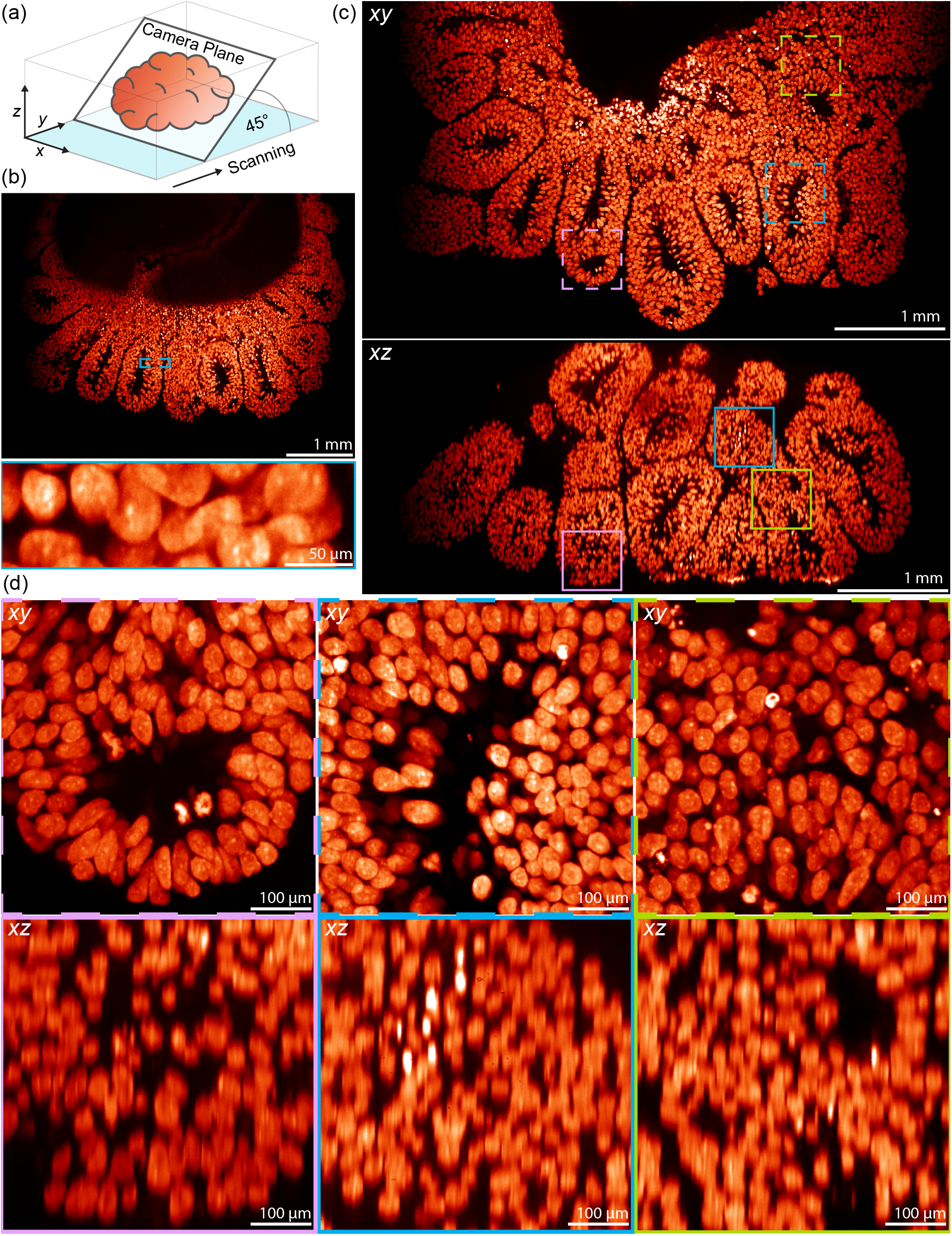
Application of DvOPM to imaging 4×-expanded brain organoids. (a) Illustrates the coordinate system used to acquire and present the data. (b) Presents an unprocessed camera frame, and a zoomed in insert, demonstrating the single frame FOV of the system. (c) Shows xy and xz slices through the reconstructed volume. (d) Gives magnified views of three areas in the xy and xz slices presented in (b). The magenta, green and cyan boxes shows the location of the respective areas in (b).

## 4 Discussion

Here we present a simple and cost-effective method for mesoscopic oblique plane microscopy. By detecting fluorescence with a camera in the remote space we eliminate the need for optical components to redirect the light, significantly improving photon throughput and reducing complexity. Although increasing the incident angle of light on the camera reduces detection efficiency, we demonstrate that this effect is minimal, resulting in an overall photon efficiency that surpasses all other reported mesoscopic OPM systems. Whilst the system we have presented here is for low-resolution, large-scale imaging of mesoscopic samples, it is important to highlight that the concept of DvOPM is fully applicable to high-resolution, single-cell-scale systems. The only limiting factor in this application is the availability of cameras with sufficiently small pixels to satisfy Nyquist sampling. The Samsung ISOCELL HP3 sensor already has a 560 nm pixel pitch, which would significantly enhance resolution over this work and, if cameras with pixels on the scale of ∼100 nm can be produced, DvOPM would provide a method for OPM imaging at the diffraction limit for all available NAs.

Alternatively, optical elements could be used to make subpixel shifts of the remote space image. This would allow an image with smaller pixels to be computationally reconstructed from multiple shifted images, albeit at the cost of reduced temporal resolution due to the requirement for multiple images per image plane. Additionally, axial resolution can increased through the technique of axial sweeping of the light sheet [10, 18].

In the implementation presented in this work, temporal resolution is limited by sample scanning via a motorised translation stage. However, high temporal resolution mirror-based scanning methods used in other mesoOPM systems could be used instead. Overall, DvOPM provides the simplest, most efficient optical arrangement to image mesoscopic biological samples in an OPM format. Furthermore, continued development of small pixel camera technologies shows exciting potential for DvOPM to be applied to higher resolution applications.

## 5 Acknowledgements

JRL and JDM thank Taylors of Harrogate for producing the calming beverage Yorkshire Tea, consumed after sensor cover glass removal. This work was funded through a Royal Society University Research Fellowship to JDM (URF\R1\221086 & RF\ERE\221078).

