## Supplementary Material for "Direct-view oblique plane microscopy"

### A Cerebral organoid generation

An hESCs cell line (H9, female), cultured in feeder free conditions, was used to generate cerebral organoids, using a protocol modified from Lancaster et al. [1]. Cells were grown in StemFlex (Thermo Fisher, A3349401) on Matrigel growth factor reduced (8.7 µg/cm$^2$, Corning, 356230) coated plates at 37 °C with controlled 5 % CO2 and passaged every 3–4 days using 0.7 mM EDTA.

The STEMdiff™ Cerebral Organoid Kit (StemCell Technologies 08570) was used to generate optimised cerebral organoids. Single cell suspension was obtained through resuspension of stem cells in Accutase (Sigma-Aldrich, A6964). 2000 cells per well were seeded in U-bottom ultralow attachment 96 well plates (Corning, CLS7007) with EB media supplemented with 50 µM ROCK inhibitor Y27632 (Millipore, SCM075). At day 5 EBs were transitioned to neural induction media. At day 7 EBs were transferred to 6 cm dishes and incubated in expansion media with dissolved Matrigel (Corning, 356234) in a dilution of 1 to 50. After another three days of culture, organoids were incubated with Improved Differentiation Media without vitamin A (50 % (v/v) DMEM F12 (Thermo Fisher Scientific, 11330032), 50 % (v/v) Neurobasal (Invitrogen, 21103049), 1:200 (v/v) N2 supplement (Thermo Fisher Scientific 17502048), 1:50 (v/v) B27-A (Thermo Fisher Scientific, 12587010), 1:100 (v/v) GlutaMAX, 1:200 (v/v) MEM-NEAA, 2.5 µg/mL insulin solution (Sigma-Aldrich, I9278), 50 µM $\beta$-mercaptoethanol (Life Technologies, 31350-010), 1:100 (v/v) penicillin-streptomycin).

### B Expansion Microscopy

Cerebral organoids were fixed for 1 hour using 4 % PFA at room temperature. After fixation, organoids were washed in 1× PBS before marination overnight in a solution of 1.4 % formaldehyde (28906, Fisher Scientific) and 2 % acrylamide (A9099, Sigma). Samples were then embedded in monomer solution (19 % sodium acrylate (Sc-236893, Santa Cruz Biotechnology), 10 % acrylamide, 0.1 % BIS (J66710.30, Thermo) in 1× PBS) supplemented with TEMED (411019, Sigma) and APS (A3678, Sigma) to a final concentration of 0.5 % and marinated at 4 °C for 2 hours. The humidified chamber containing the embedded samples was then heated to 37 °C for 1.5 hours. The organoid-containing gels were then incubated with denaturation buffer (200 mM SDS (30018385, Fischer Scientific), 200 mM NaCl (3624-01, J. T. Baker), 50 mM Tris (30018880, Fischer Scientific), pH 9) for 15 minutes at room temperature followed by 1 hour and 30 minutes at 95 °C. Gels were then subject to a first round of expansion by incubating three times in dd$H_2O$ for 30 minutes. The gel was then shrunk by washing 3 times with PBS for 15 minutes. To label nuclei, gels were incubated with Hoechst 33342 diluted in PBS-BSA 2 % for 48 hours. Gels were then washed in PBS-Tween 0.1 % (P7949, Sigma) and subject to a final round of expansion by incubating three times in dd$H_2O$ for 30 minutes.

### C Removal of camera sensor cover glass

The Ximea MU196MR-ON camera used in this work is equipped with an Onsemi AR2020 sensor, which is manufactured with a 550 µm sensor cover glass as standard. However, as the camera must be tilted in an image plane, the presence of this glass introduces significant astigmatism in the final images, as shown in Fig. S2 and S3. To eliminate this optical aberration and achieve optimal imaging performance, the sensor cover glass needed to be removed.

The removal process began with the careful disassembly of the camera to expose the sensor and provide unobstructed access. Once the sensor was accessible, the remaining steps were performed under a stereomicroscope. Using a sharp pair of tweezers, the adhesive securing the edges of the sensor cover glass to the plastic sensor mount was delicately removed. After the adhesive at the edges was cleared, the serrated tip of the tweezers was used to cut away a corner of the plastic sensor mount, allowing access to the underside of the glass window. A razor blade was then inserted between the bottom of the sensor glass and the plastic mount. The adhesive bonding the glass to the mount was carefully loosened and scraped away using the blade, which was then used as a lever to gently separate the glass from the plastic mount.

This process was repeated incrementally around the perimeter of the glass, with the razor blade repositioned to ensure uniform separation until the adhesive bond was completely broken and the glass became fully detached. Once the glass was freed, the sensor was inverted to allow the glass to fall away from the sensor surface. A rubber bulb blower was subsequently used to remove as many fine glass fragments as possible, while taking great care not to damage the exposed gold contacts on the sensor. After the removal process was completed, the camera was reassembled and mounted in the DvOPM system using a custom 3D-printed mount, which offered additional protection against dust contamination.

The procedure successfully removed the sensor cover glass without any damage to the camera pixels. However, despite efforts to perform the process as cleanly as possible, a small number of glass fragments adhered to the sensor surface creating small shadow artefacts in the image, as seen in the lower middle image of Fig. 3c. However, the vast majority of the sensor surface remained clean and unaffected.

While this work demonstrates that manual removal of the sensor cover glass is feasible without requiring specialised equipment or expertise, it is recommended that this procedure be performed professionally. Sensor cover glass removal is a standard practice for enhancing both the sensitivity and spectral range of cameras used in optical applications and as such several companies offer professional services for this process to minimise the risk of contamination or sensor damage.

### D DvOPM optimised for different clearing methods

The Nyquist-Shannon sampled pixel limited resolution in DvOPM is twice the effective size of the camera pixels in the sample. Therefore, lateral imaging resolution is limited by the combination of the camera pixel size and the magnification of the remote refocus system.

Camera pixel size is primarily limited by the smallest commercially available camera pixels. Therefore, with future development of smaller camera pixels, the camera pixel size can be decreased arbitrarily. Conversely, whilst the remote refocus magnification can theoretically be tuned to anything, to satisfy the aberration free imaging condition, as is required to image large volumes, it must be equal to the refractive index of the sample. Therefore, to obtain the best imaging performance, the DvOPM system should be tuned to match the refractive index of the intended sample. In the case of this work, the intended sample is 4×-expanded brain organoids and as such the magnification of remote refocus is set to 1.33.

When applying DvOPM to cleared tissues, since the method for clearing affects the final refractive index, it is important to design the system for the intended clearing method. Additionally, this means that, for a given camera, the final pixel limited lateral resolution is dependant on the clearing method. Clearing methods with a higher refractive index need a higher magnification remote refocus, which then makes the effective size of the pixels in the sample smaller and thus improves the pixel limited resolution.

Table S1 compares the achievable pixel limited lateral resolution for DvOPM systems designed to image samples prepared using common optical clearing methods [2]. We present the resolution for DvOPM systems constructed using three different camera sensors: an Onsemi AR1630 with 1 µm pixels (smaller pixels but currently only available as RGB), an Onsemi AR202 with 1.4 µm pixels (used in this work) and a Sony IMX226 with 1.85 µm pixels (available from Allied Vision without a sensor cover glass). It can be seen that by using higher refractive index clearing method, such as BABB or iDISCO, and using the Onsemi AR1630 sensor, the pixel limited lateral resolution can be improved by 40 %, from 2.11 µm to 1.28 µm, compared to this work.

**Supplementary Table 1:** Comparison of pixel limed lateral resolution for different optical clearing methods.

| **Clearing protocol** | **Ref** | **RI** | **Pixel limited lateral resolution / µm**<br>**Onsemi AR1630**<br>1 µm pixels | **Onsemi AR2020**<br>1.4 µm pixels | **Sony IMX226**<br>1.85 µm pixels |
|---|---|---|---|---|---|
| *Solvent-based tissue clearing techniques* | | | | | |
| BABB | [3] | 1.56 | 1.28 | 1.79 | 2.37 |
| Fluoclear BABB | [4] | 1.55–1.56 | 1.28–1.29 | 1.79–1.81 | 2.37–2.39 |
| Modified BABB | [5] | 1.55–1.56 | 1.28–1.29 | 1.79–1.81 | 2.37–2.39 |
| 3DISCO | [6] | 1.56 | 1.28 | 1.79 | 2.37 |
| iDISCO | [7] | 1.56 | 1.28 | 1.79 | 2.37 |
| uDISCO | [8] | 1.55–1.57 | 1.27–1.29 | 1.78–1.81 | 2.36–2.39 |
| a-uDISCO | [9] | 1.55–1.57 | 1.27–1.29 | 1.78–1.81 | 2.36–2.39 |
| fDISCO | [10] | 1.56 | 1.28 | 1.79 | 2.37 |
| vDISCO | [11] | 1.55–1.57 | 1.27–1.29 | 1.78–1.81 | 2.36–2.39 |
| Ethanol-ECi | [12] | 1.56 | 1.28 | 1.79 | 2.37 |
| 2ECi | [13] | 1.56 | 1.28 | 1.79 | 2.37 |
| PEGASOS | [14] | 1.54 | 1.30 | 1.82 | 2.40 |
| SHANEL | [15] | 1.55–1.56 | 1.28–1.29 | 1.79–1.81 | 2.37–2.39 |
| FOCM | [16] | 1.50 | 1.33 | 1.87 | 2.47 |
| *Aqueous-based tissue-clearing techniques* | | | | | |
| SeeDB | [17] | 1.49–1.50 | 1.33–1.34 | 1.87–1.88 | 2.47–2.48 |
| FRUIT | [18] | 1.50 | 1.33 | 1.87 | 2.47 |
| TDE | [19] | 1.45 | 1.38 | 1.93 | 2.55 |
| ScaleS | [20] | 1.44 | 1.39 | 1.94 | 2.57 |
| CUBIC-X | [21] | 1.47 | 1.36 | 1.90 | 2.51 |
| CUBIC-P | [22] | 1.52 | 1.32 | 1.84 | 2.43 |
| UBasM | [23] | 1.47–1.48 | 1.35–1.36 | 1.89–1.90 | 2.5–2.51 |
| *Hydrogel embedding-based tissue-clearing techniques* | | | | | |
| CLARITY | [24] | 1.45 | 1.38 | 1.93 | 2.55 |
| PACT | [25] | 1.44 | 1.39 | 1.94 | 2.57 |
| PARS | [25] | 1.45 | 1.38 | 1.93 | 2.55 |
| ACT-PRESTO | [26] | 1.45 | 1.38 | 1.93 | 2.55 |
| Bone CLARITY | [27] | 1.47 | 1.36 | 1.90 | 2.51 |
| SWITCH | [28] | 1.46 | 1.37 | 1.92 | 2.53 |
| SHIELD | [29] | 1.46 | 1.37 | 1.92 | 2.53 |
| **ExM** | **[30]** | **1.33** | 1.50 | **2.11** | 2.78 |

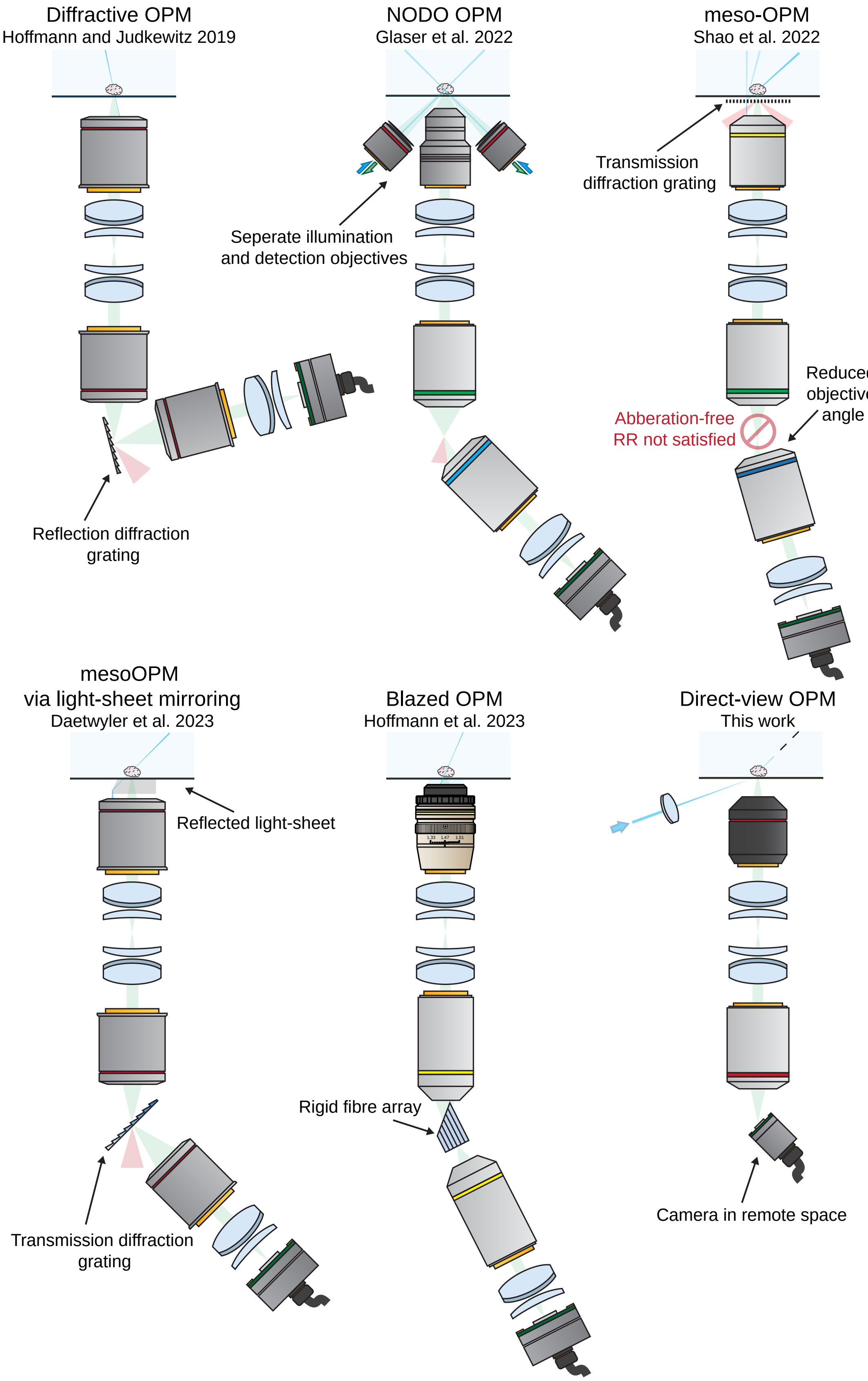


**Supplementary Figure 1:** Comparison of existing mesoscopic OPM systems. In each diagram the excitation light is show in blue, the collected emission light is shown in green and the uncollected emission light, due to system inefficiencies, is shown in red. Additionally, the main components enabling the technique are labelled.

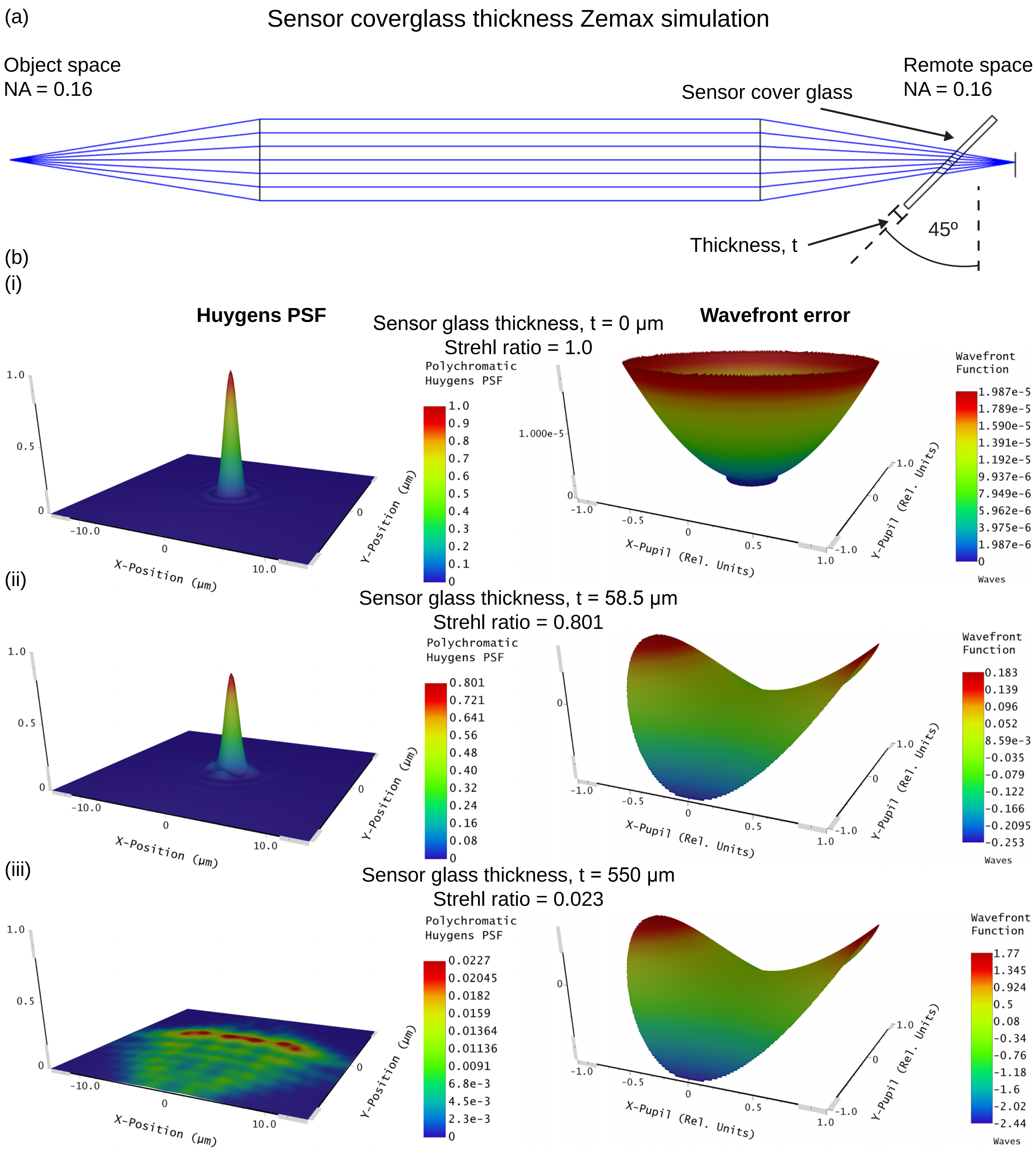


**Supplementary Figure 2:** Zemax simulation demonstrating the effect of imaging through varying thicknesses of glass at 45° for a remote refocus system with unity magnification and an NA of 0.16. This demonstrates the aberrations introduced by the sensor cover glass when imaging at 45° highlighting the requirement for the removal of this glass window. (a) Shows an diagram of the optical system used for the Zemax simulation. (b) Shows both the Huygens PSF, Strehl ratio and wavefront error when imaging through glass of a thickness of 0, 58.5 (limit of diffraction limited imaging) and 550 μm. Note that the Onsemi AR2020 sensor used in this work is manufactured with a 550 μm thick cover glass.

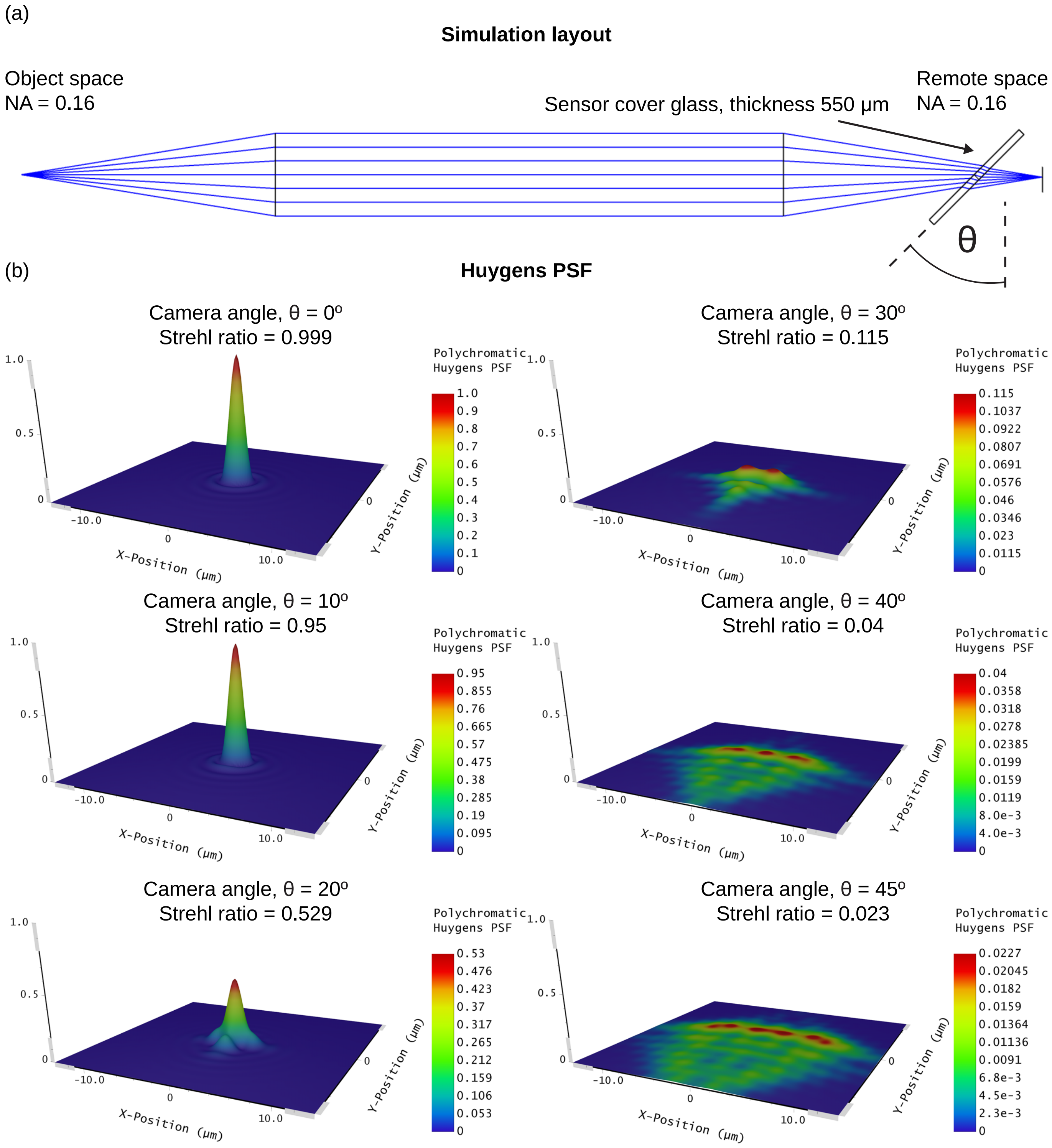


**Supplementary Figure 3:** Zemax simulation demonstrating the effect of imaging through a 550 µm thick glass window at varying angle for a remote refocus system with unity magnification and an NA of 0.16. This demonstrates the aberrations introduced by the sensor cover glass when imaging at 45° highlighting the requirement for the removal of this glass window. (a) Shows an diagram of the optical system used for the Zemax simulation. (b) Shows both the Huygens PSF and Strehl ratio when imaging through glass of a thickness of 550 µm and an angle of 0, 10, 20, 30, 40 and 45°.

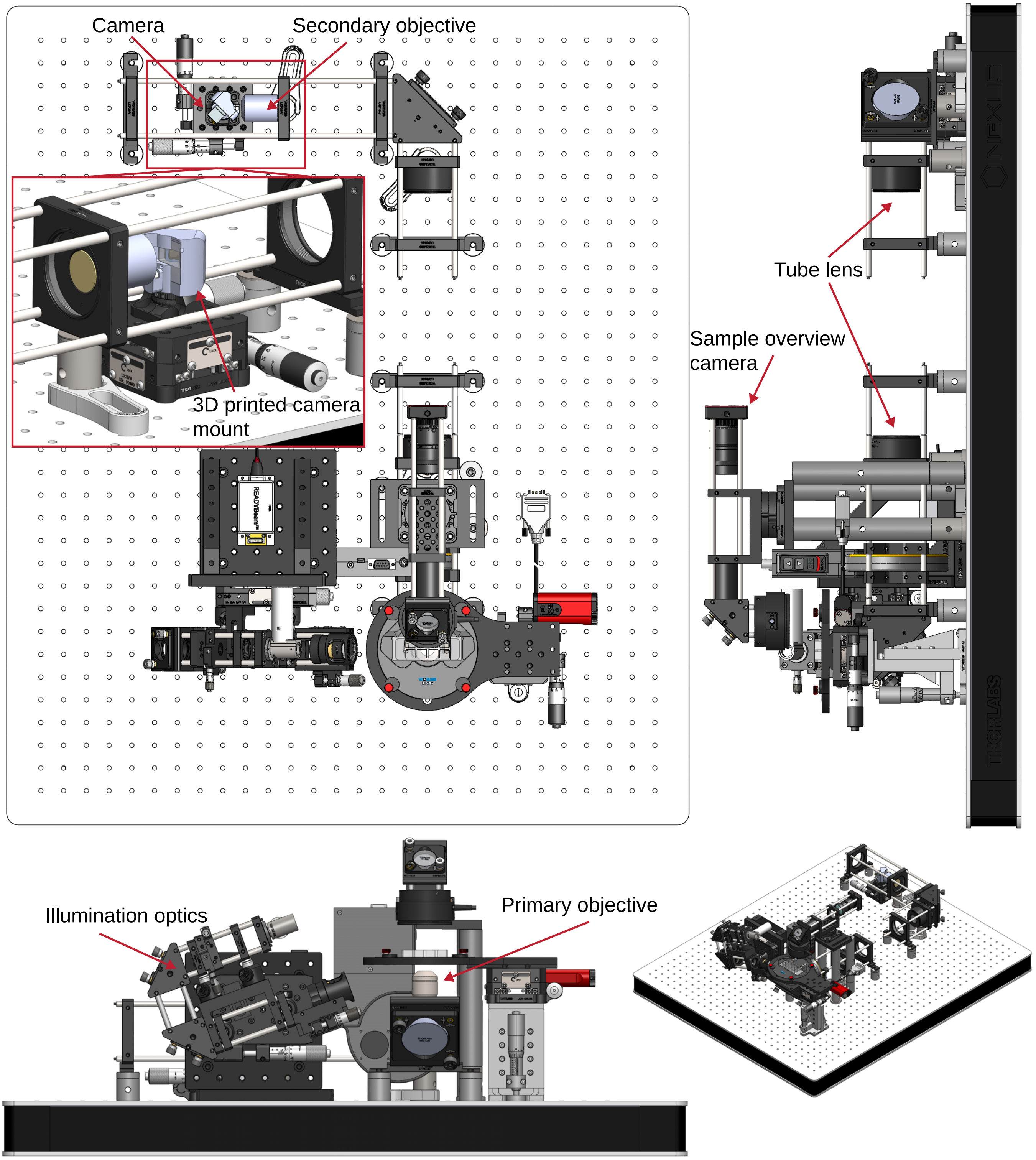


**Supplementary Figure 4:** CAD rendering of the DvOPM system constructed in this work. The important components in the system are highlighted with labels and magnified view of the camera held in the custom 3D printed mount is presented in the red box. It should be noted that, with the exception of the two objective lenses, laser source and camera, all the components used to build the system presented here are standard Thorlabs components.

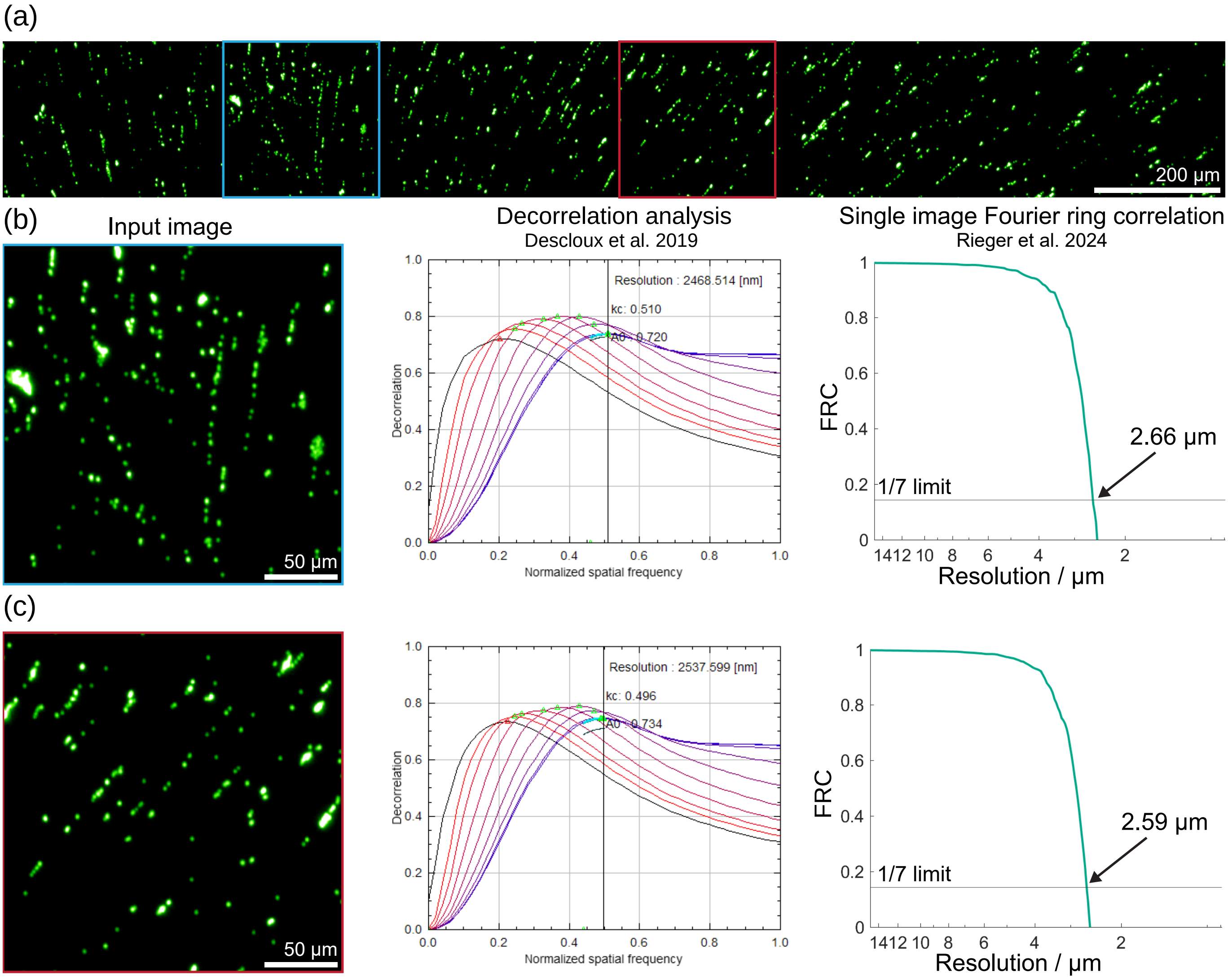


**Supplementary Figure 5:** Measurement of the lateral spatial resolution calculated using image decorrelation analysis [31] and single image FRC [32]. The analysis was performed on a single XY slice of 200 nm beads dried onto a coverglass used for the FWHM measurement presented in the main text (shown in a). Panels (b) and (c) show plots of the decorrelation analysis and single image FRC when performed on two square regions of the FOV shown in (a). Decorrelation analysis was performed using the ImageJ image decorrelation analysis plugin provided by Descloux et al. [31] and the single image FRC was perfromed using the Matlab script provided by Rieger et al. [32].

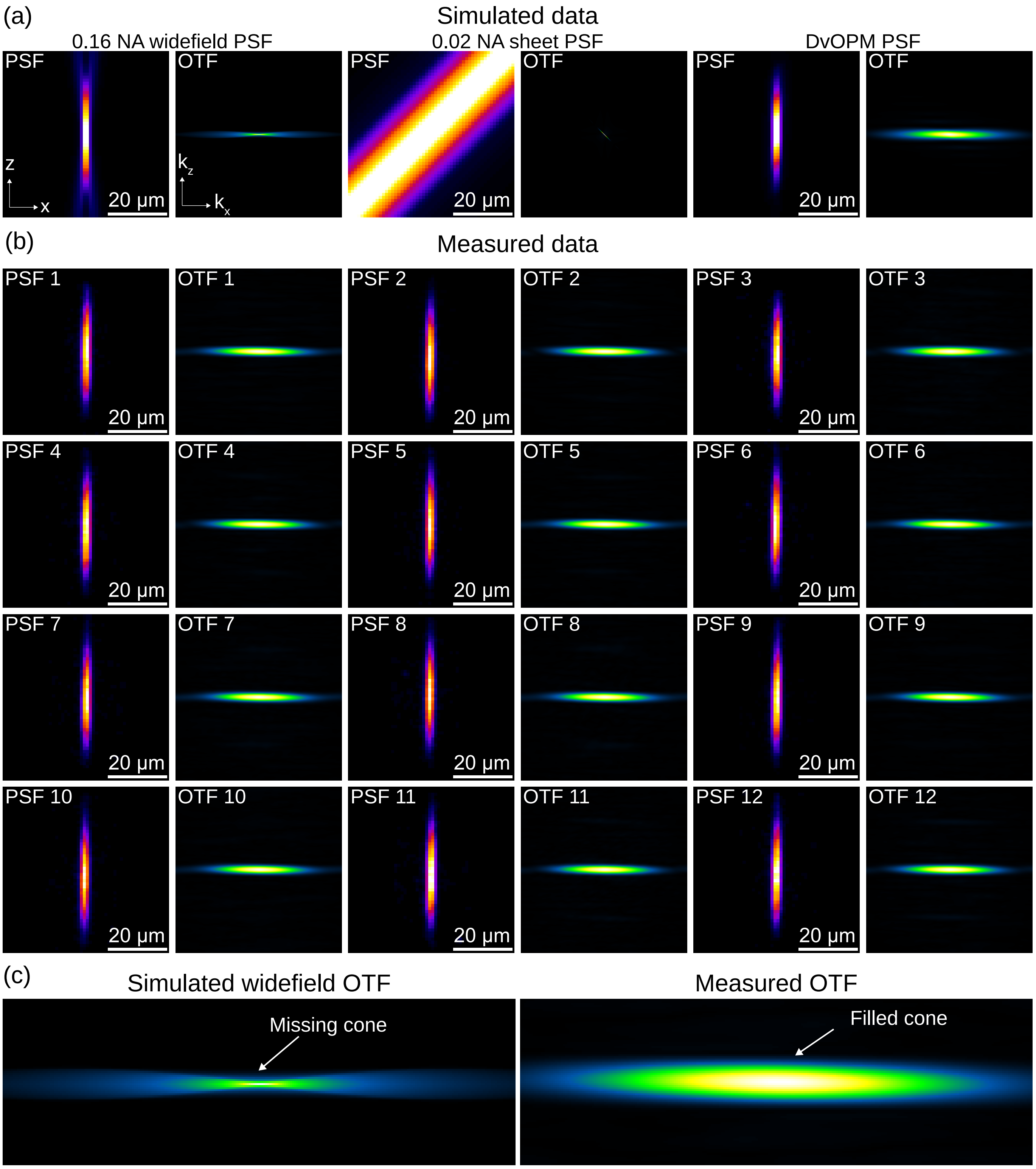


**Supplementary Figure 6:** Demonstration of DvOPM optical sectioning via comparison of simulated widefield PSF and OTF to the measured PSF and OTF for the system. (a) presents simulated results for the PSF and OTF for a 0.16 NA widefield, 0.02 NA sheet and final DvOPM. The DvOPM PSF is calculated as the product of the widefield and sheet PSF and the DvOPM OTF is the convolution of the widefield and sheet OTF. (b) shows the measured PSF and OTF from 12 different 200 nm beads. (c) compares the simulated widefield OTF to the measured DvOPM OTF highlighting how the missing cone is filled, thus achieving optical sectioning.

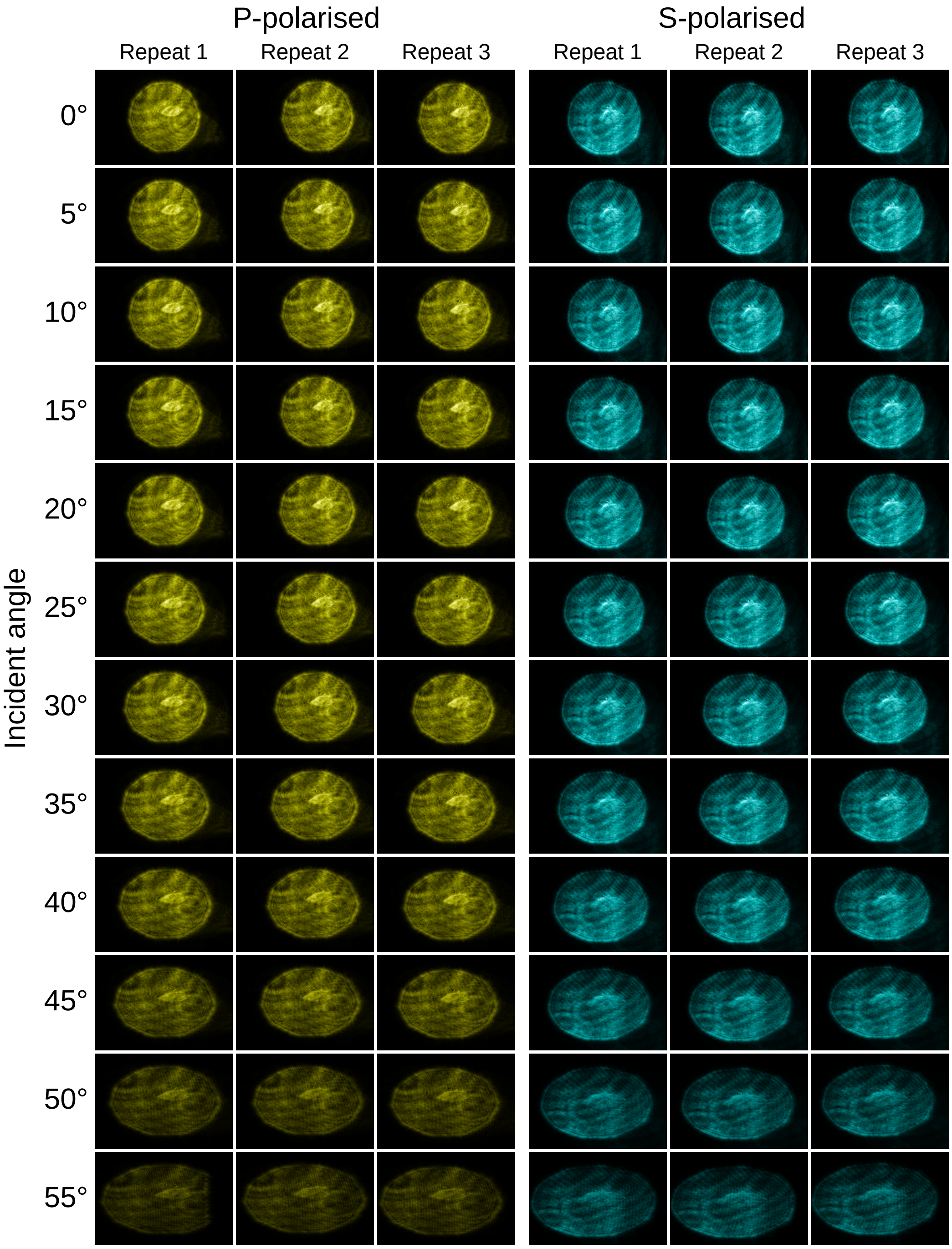


**Supplementary Figure 7:** Raw data from the camera efficiency with incident ray angle investigation. A collimated beam of both P (yellow) and S (blue) polarisation is incident on the camera at an angle ranging from 0 to 55° in 5° increments. Note that the images shown in this figure have been cropped down from the original data for display purposes.
